# Changes in the three-dimensional microscale topography of human skin with aging impact its mechanical and tribological behavior

**DOI:** 10.1101/2020.10.18.344606

**Authors:** Juan G. Diosa, Ricardo Moreno, Edwin L. Chica, Junes A. Villarraga, Adrian Buganza-Tepole

## Abstract

Human skin enables interaction with diverse materials every day and at all times. The ability to grasp objects, feel textures, and perceive the environment depends on the mechanical behavior, complex structure, and microscale topography of human skin. At the same time, abrasive interactions, such as sometimes occur with prostheses or textiles, can damage the skin and impair its function. Previous theoretical and computational efforts have shown that skin’s surface topography or microrelief, is crucial for its tribological behavior. However, current understanding is limited to adult surface profiles and simplified two-dimensional simulations. Yet, the skin has a rich set of features in three dimensions, and the geometry of skin is known to change with aging. Here we create a numerical model of a dynamic indentation test to elucidate the effect of changes in microscale topography with aging on the skin’s response under indentation and sliding contact with a spherical indenter. We create three different microrelief geometries representative of different ages based on experimental reports from the literature. We perform the indentation and sliding steps, and calculate the normal and tangential forces on the indenter as it moves in three distinct directions based on the characteristic skin lines. The model also evaluates the effect of varying the material parameters. Our results show that the microscale topography of the skin in three dimensions, together with the mechanical behavior of the skin layers, lead to distinctive trends on the stress and strain distribution. The major finding is the increasing role of anisotropy which emerges from the geometric changes seen with aging.

## Introduction

Our perpetual interaction with the environment is mediated by the skin, the largest organ in our body [1, 2]. Contact with the materials that surround us enable us to understand and interpret the things we touch, from perceiving shape and texture, to manipulating objects dexterously [3, 4, 5, 6]. Unfortunately, mechanical interaction between the skin and the environment can also lead to injury, from superficial abrasion to chronic wounds [7, 8, 9]. This is especially true in situations in which there is constant contact, cyclic relative motion, and large tangential and normal forces applied to the skin [10, 11]. A primary example of this kind of scenario is the interaction between skin and prostheses [12, 13, 14, 15]. Another example is pressure sore initiation in individuals who are bed- or wheelchair-ridden [16, 17, 18]. Among other factors, such as ischemia and inflammation [19, 20], friction between the skin of these individuals and textiles or common materials used in wheelchair or prostheses design can contribute to abrasion and skin damage, paving the way for more serious chronic wounds [7, 21, 22, 23]. Other applications for which anticipating skin pressure and friction are important include the design of shoes and clothing for diabetic individuals, wearable robots for rehabilitation, exoskeletons for soldiers, and sports equipment [24].

Skin has remarkable mechanical properties that enable its function. This organ is strong and tough yet flexible. It operates physiologically in the large deformation regime and shows a highly nonlinear stress-strain response. The frictional behavior of human skin at the tissue scale (in the order of cm) is affected by several variables including age [25, 26], anatomical region [27, 28, 29], contact material [30], type of contact [5, 31], environmental conditions [6, 26, 28], and wetness in the tissue [32, 16]. However, to discriminate between the relative contributions of these factors, it is advantageous to zoom in to the mesoscopic scale, in the order of hundreds of *μ*m to a few mm. At this scale, the friction between the skin and a given material depends strongly on the inherent mechanical features seen at this scale: the mechanical properties of the different skin layers, the deformation of the tissue during contact, and the topography of the skin, also referred to as microrelief [33, 34]. However, our knowledge of skin friction at this mesoscopic scale has several, important gaps. In particular, here we are chiefly interested in elucidating how changes of skin threedimensional (3D) structure with aging may alter the mechanical and frictional behavior of this tissue.

For such a thin structure, the skin has a fascinating architecture. The tissue can be divided into three layers. At the top there is the epidermis, below that the dermis, and the hypodermis is the bottom layer, connecting the skin to the underlying muscle [36, 37]. The sublayer of the epidermis right above the interface with the dermis is populated by keratinocyte cells. As these cells proliferate and differentiate they move up through the epidermis. Some of the changes in keratinocyte cells associated with differentiation include accumulation of keratin, gradual disintegration of organelles, and eventually cell death [38, 39]. In fact, the outermost sublayer of the epidermis, the stratum corneum (SC), is made out of dead cells called corneocytes, which are constantly released to the environment via desquamation [40]. The biological changes during cell differentiation in the epidermis have corresponding changes in mechanical properties. Commonly, two different mechanical responses are distinguished for the top layer of the skin, that of the living part of the epidermis, and that of the SC. The middle layer of the skin, the dermis, has a completely different structure compared to the epidermis and, consequently, also shows unique mechanical function. The dermis is a collagen-based connective tissue that is mainly responsible for the tensile properties of skin [41]. Underneath the dermis, the hypodermis or subcutaneous tissue has yet a different composition and mechanical behavior. The hypodermis is made out of fat cells and tissue, serving as a loose viscoelastic attachment to the underlying muscle. Remarkably, the complex skin microstructure is packed in a thickness of only 1-3mm (see Figure 1).

**Figure 1:**
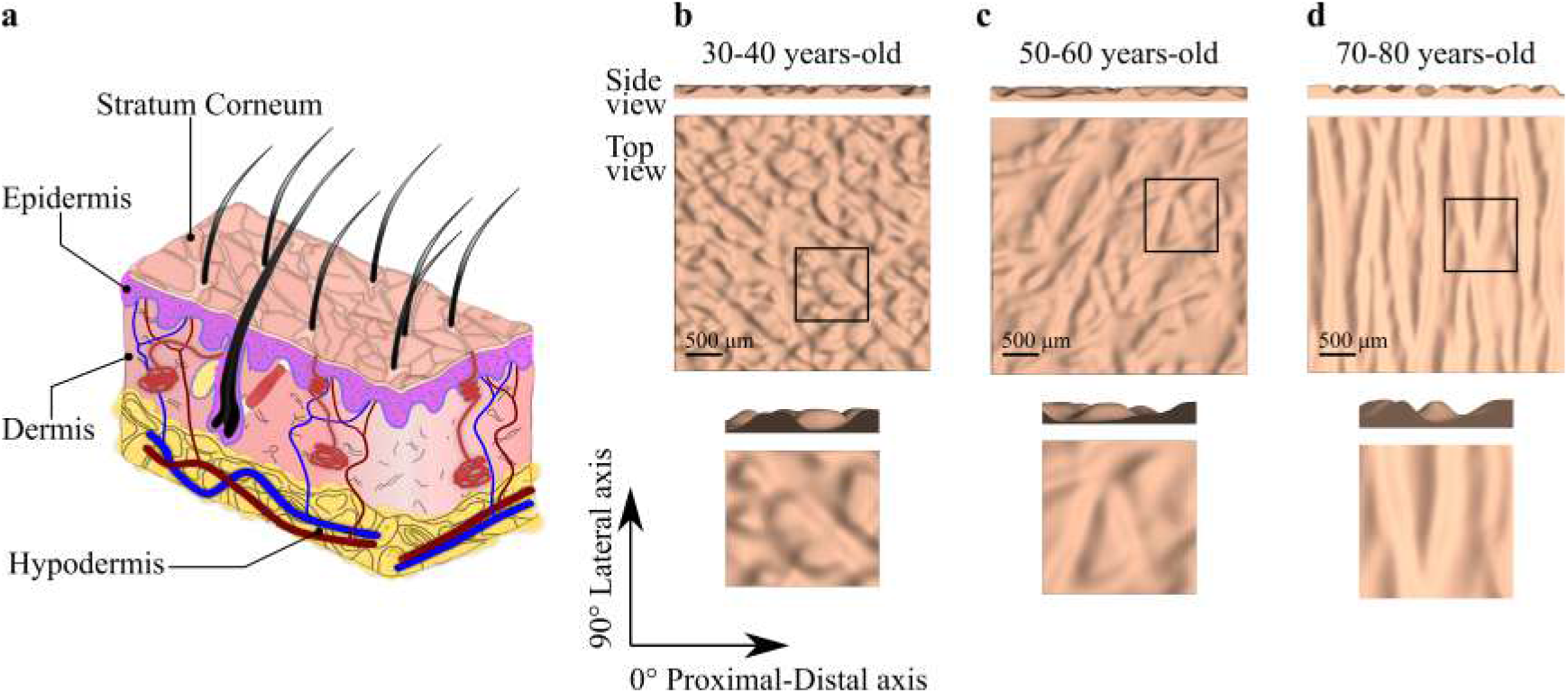
Human skin anatomy and topography: **a**) Schematic of human skin layers, **b**) 30 – 40 years-old topography, **c**) 50 – 60 years-old topography and **d**) 70 – 80 years-old topography. Three-dimensional surface geometries are reconstructed based on the work by Zahonuani et al [35]

In addition to its organization across the thickness, the top surface and the interfaces between the skin layers also show a non-trivial geometry. The interface between the epidermis and the dermis is a two-dimensional surface with a wave pattern of peaks and valleys called the rete ridges [42, 43, 44]. The outermost surface of the skin also shows an intricate microscale topography or microrelief. This topography is usually characterized by furrows on the surface of the skin. The shape and characteristics of these furrows are delimited by different lines classified by their wideness and deepness into primary, secondary, and in some cases even tertiary and quaternary lines [44, 45, 46]. The primary lines are the deepest and describe the contours of the visible polygons and plateaus on the skin surface. Primary lines help to control sebum quantity and movement of sweat. The secondary lines go across the plateaus formed by the primary lines, connecting the adjacent plateaus and furrows. Finally, the tertiary and quaternary are delimited by the corneocytes borders and surface characteristics of the corneocytes respectively [47, 48]. The skin surface, mainly the area of the plateaus and the direction of the primary lines have been identified as key for the frictional response [49, 50].

The skin, like most biological tissues, shows inherent variability in structure and mechanical properties within and across individuals. For instance, even focusing on the skin of an individual, there is variation with respect to anatomical location and the environment conditions. Hydration and swelling of the corneocytes through water absorption [51] leads to reduction of the apparent stiffness of the SC by orders of magnitude compared to the baseline physiological state [40, 52]. Those changes in the SC mechanics are reflected in a broad range of values for the friction coefficient between skin and different materials [25, 30, 53, 54]. For example Klaassen et. al. have found that a change in relative humidity from 40% to 80% leads to up to a twofold increase in the friction forces [55, 53].

Also common across living tissues, skin does not remain unchanged throughout our lifetime, but it adapts its structure and mechanical behavior based on sun exposure, mechanical cues, disease, and aging [45, 46, 47]. In particular, dramatic changes of skin microrelief characteristics with aging have been reported (see Figure 1) [45, 35]. The main trends are changes in the direction, depth and width of the primary lines [46, 35, 56]. In consequence the density of the furrows decreases, incrementing the plateaus area [57, 58]. Changes in morphology, composition, and mechanical properties have also been reported with aging. For instance, the dermis becomes thinner, the tissue has more slack or becomes more compliant under small deformations, but is stiffer at larger stretches [59, 60].

Experimentally, the coefficient of friction for human skin in contact with different materials is usually evaluated as the ratio between the recorded tangential and normal forces. These measurements have been almost entirely restricted to the macroscopic scale (on the order of cm). The results from such experiments have shown that the frictional response of human skin at this scale cannot be accurately described by common models such as Amontons – Coulombs [3, 5, 61, 62]. Several other, more complex descriptions have been proposed, including statistics regression models [25, 26], power laws [3, 28, 38, 54], adhesion models [63] and others [64]. These efforts are focused on capturing the macroscopic scale phenomena. Yet, a major gap in our knowledge is the limited understanding of the role played by the mesoscale structure and mechanical response in producing the observed friction at the larger spatial scales. It is extremely difficult to experimentally dissect the fundamental mechanisms occurring at the mesoscale because of the inherent challenge in measuring the force and contact area distributions with a fine enough resolution in relevant scenarios. Instead, high-fidelity computational models can help bridge this gap. For example, Leyva-Mendivil et. al. [33, 34] used a two-dimensional model of skin to analyze the influence of the rigidity of the SC, indenter radius, indentation depth and local coefficient of friction in the macroscale friction behavior of skin. The results of these studies show the importance of the skin surface topography and SC stiffness in the deformation component of the macroscopic friction. Yet, even though the surface of the skin has a rich set of features in 3D, and the geometry of skin is known to change with aging, the effect of these features on the mechanical and tribological response of skin has not been thoroughly investigated.

In this paper we build a detailed 3D finite element model of skin in contact with a spherical indenter in order to elucidate the effect of microrelief changes with aging. Realistic computational models of skin mechanics are now possible due to the comprehensive characterization and constitutive modeling of human skin over the past few decades. Skin has been studied with uniaxial, biaxial [65, 66], multiaxial [67], bulge [68], suction [69], and indentation tests. Several constitutive models have been proposed pertinent to different applications, loading conditions, and time scales. For example, linear [70], viscoelastic [71, 72], multiphasic [1], and hyperelastic [73, 74] constitutive models have been used. There have also been efforts to distinguish between the mechanical behavior of the different skin layers [75]. In fact, the influence of the different mechanical properties of individual layers is more evident in scenarios where the skin is under compression [76, 77].

## Methods

We create detailed 3D finite element models of skin based on reported measurements of the skin microscale topography available in the literature [35, 50, 57, 78]. In particular, the work by Zahouani et. al. has led to a thorough characterization of the changes in skin microrelief with aging, with geometric features on the order of 160 *μ*m. We use their data to reconstruct surfaces in 3D, and then extrude these surfaces computationally to get the solid 3D geometry.

### Reconstruction of skin surface geometries

We base our models on the 3D confocal microscope analysis of human skin topography by Zahouani et al [35]. The z-height contours of the left forearm skin of 3 representative subjects were processed, one contour from each of the following age groups: 30 – 40, 50 – 60, and 70 – 80 years-old. A section of 3.5 mm x 3.5 mm from each image is selected and pre-processed on GIMP 2.10. To deal with the excessive noise in the contour images, which would result in unrealistic surface profiles, we apply a 5×5 pixel Gaussian blur filter and scale all the images to 100 x 100 pixels using a cubic interpolation method. A Matlab ^®^ (version R2018a; The MathWorks Inc., Massachusetts, USA) script then reads from the PNG file the pixel color and assigns a height value based on the contour scale from the original images [35]. The surfaces built in this manner are further smoothed out by applying a Gaussian filter with *σ* =0.75 pixels to relocate the remaining extreme outlier heights. The surfaces capture the distinct geometric features of different ages. We analyze the topography of each age group using ImageJ 1.52a to determine the direction of the primary skin lines. In Figure 1, the 0° direction is aligned with the proximal-distal axis of the arm, from the elbow to the wrist, and the 90° direction is aligned with the lateral axis. Two different quadrants are analyzed, the first one between 0° to 90° and the second one between 90° and 180°. Primary lines in the images are identified manually in both quadrants separately, the direction of these lines is saved, and the median from five measurements for each age are computed. In the 30-40 years-old geometry, the primary lines show orientations 43.3° and 136.5°; in the 50-60 years old case the lines are oriented at 46.3° and 103.9°; and in the 70-80 years-old skin the primary lines are oriented at 81.4° and 100.4°.

### Generation of 3D finite element meshes

We create a finite element mesh for each of the geometries with a custom Matlab script. The mesh has 12 elements through the thickness, with element-element interfaces placed so as to capture four anatomical layers: stratum corneum (SC), viable epidermis, dermis and hypodermis, with thicknesses 0.02mm [13, 52, 79], 0.05 mm [76], 0.84 mm [76] and 2 mm [80], respectively. Analogous to the top surface topography, the epidermis-dermis interface shows a similar profile [37, 79, 81], but the subsequent interfaces between skin layers can be assumed almost flat [33, 45, 76]. Hence, we apply linear smoothing of the surface profile as we extrude the mesh through the thickness. To avoid boundary effects in the simulations of the skin in contact with the spherical indenter, we embed the mesh generated with our Matlab code into a larger domain (Figure 2a). The extension of the skin model is created on preview 2.1 (MRL The University of Utah, MBL Columbia University). This additional portion of the domain also has four anatomical layers with different properties, but lacks the surface topography. To match the nodes at the interfaces of both meshes, a transition region is also prescribed in the Matlab-generated mesh. The indenter used in this study is a smooth hemisphere of 5mm diameter. The models created have an average of 367 522 nodes and 340 065 Hex8 elements.

**Figure 2:**
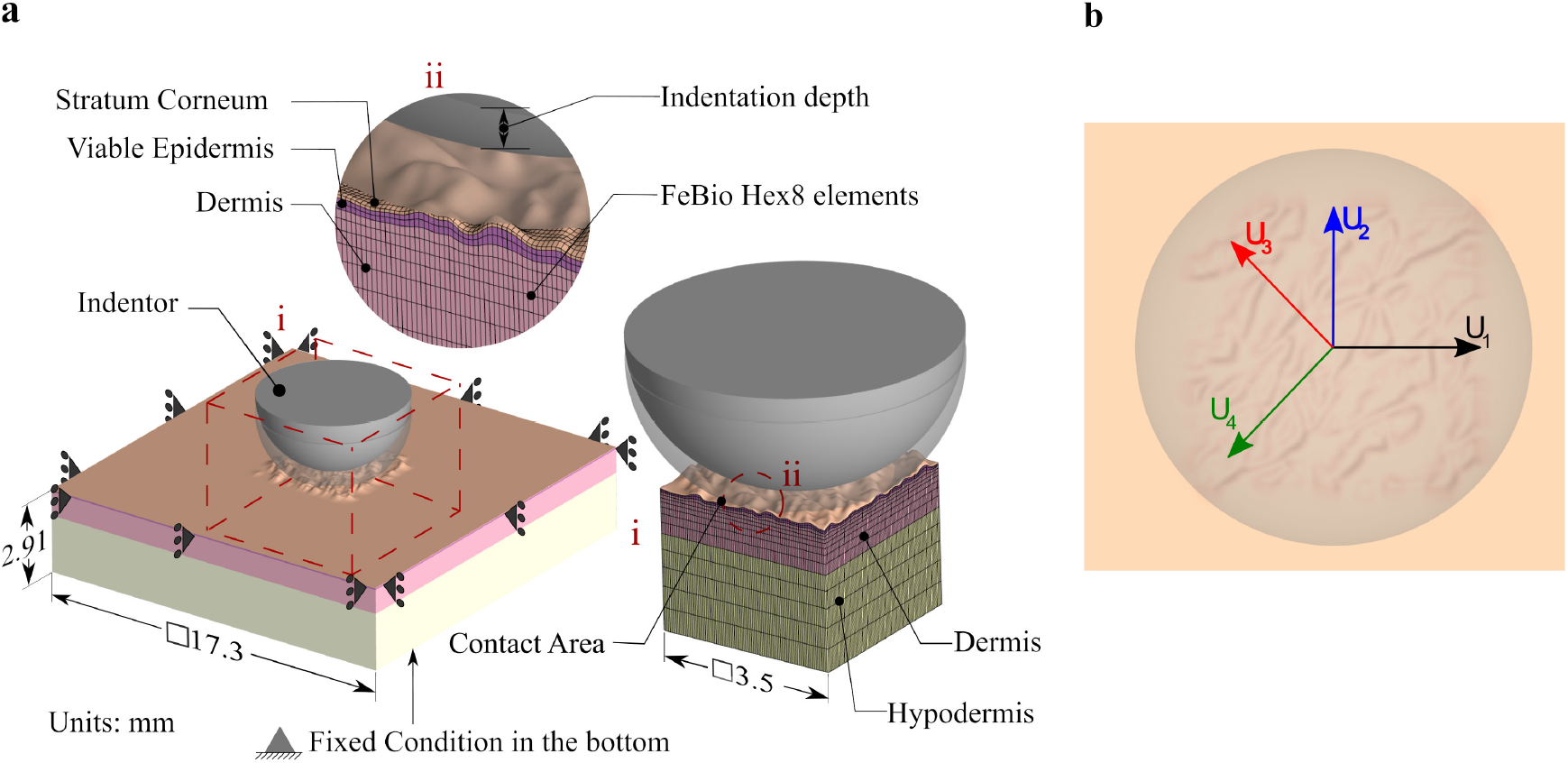
Finite element simulation setup **a**) Indentation step. The domain is more than three times larger than the indenter radius to minimize boundary effects. The spherical indenter interacts with a portion of the domain that has a detailed microrelief geometry, shown in the zoomed region (i). The mesh consists of Hex8 elements, with 12 elements through the thickness to capture the four layers of skin. Additional detail is shown in the subpanel (ii). **b**) Indenter directions of movement in displacement step are decided based on the anatomical axis (lateral *U*_1_, proximal-distal *U*_2_)as well as perpendicular to the primary skin lines (*U*_3_, *U*_4_).

### Mechanical behavior of skin

The viable epidermis, dermis and hypodermis are assumed to follow the Ogden hyperelastic constitutive relation, with material parameters obtained from the literature (Table 1) [76]. The SC is modeled as a Neo-Hookean solid. To investigate the influence of the SC hydration on its mechanical behavior, two different properties are considered. The low rigidity represents the behavior of the SC under wet conditions [33, 34, 45, 82]. In the dry condition, SC is up to sixty times stiffer than in the wet condition [45, 82]. The spherical indenter is modeled as a rigid body.

**Table 1:** Material parameters for the different human skin layers [76, 45, 33]

| Layer | Parameter 1 [MPa] | Parameter 2 [-] |
| --- | --- | --- |
| SC (Dry) | $E=360$ | $\nu=0.48$ |
| SC (Wet) | $E=6$ | $\nu=0.48$ |
| Viable Epidermis | $\mu=7.74823$ | $\alpha=2.4291$ |
| Dermis | $\mu=0.0205$ | $\alpha=3.2999$ |
| Hypodermis | $\mu=0.0094$ | $\alpha=4.0319$ |

### Age-related changes in mechanical properties and thickness

In addition to the changes in microrelief with aging, the mechanical properties and microstructure of skin also change with age. Overall there is a thickness reduction of the whole skin with aging, and in particular the dermis [44, 59]. We thus performed additional simulations in which we reduced by 50% the dermal thickness of the 50-60 and the 70-80 microrelief models. According to reports of change in dermis mechanical properties with aging [83, 84, 59, 60], which suggest that there is more slack in the dermis at small stretches but potentially stiffer response at larger stretches, we considered two additional values of the shear modulus of the dermis with respect to the parameter in Table 1. In one case we reduced the shear modulus of the dermis by half, and in the other case we considered twice the value in Table 1.

### Boundary and contact conditions

The bottom surface of the model is fixed in all directions, whereas the lateral surfaces of the model satisfy symmetry conditions in their respective planes. The movement of the indenter is divided into two steps: static indentation following by dynamic contact. We consider an indentation depth of 500 *μ*m (Figure 2a). After the tip reaches the prescribed depth, the indenter moves horizontally along one of four different directions: *U*_1_, parallel to the global *x* axis which is the proximal-distal axis of the forearm; *U*_2_, parallel to the global *y* axis which is the transverse axis; *U*_3_ or *U*_4_, normal to the direction of the primary lines determined for each age (Figure 2b). We model the contact between the indenter and the skin using the *sliding elastic contact* formulation in FeBio with the augmented Lagrangian strategy. In this study we model two cases: we perform simulations with frictionless contact as a control; we then set the local coefficient of friction to be *μ_l_*=0.2. For further control, we generate equivalent models but without the microrelief, i.e. flat geometries. We compute the coefficient of friction as the ratio between the tangential forces, *f_t_* and *f_b_* (collinear and orthogonal to the indenter movement respectively), and normal forces, *f_n_* [85],

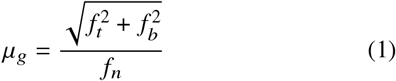

### Multivariate regression analysis

In order to quantify the influence of each factor from the simulation on the resulting coefficient of friction, we do a multivariate analysis based on generalized linear regression using R 3.6.1 (R Foundation). The parameters are codified as follows: for the age group we use 1 for the 30-40 year-old geometry, 2 for 50-60 year-old case, and 3 for the 70-80 year-old geometry; for the material properties of the skin, 1 codifies wet and 2 is used for dry; finally, the codes for the different directions are 1, 2, 3 and 4 for *U*_1_ *U*_2_, *U*_3_ and *U*_4_ respectively. The regression is thus summarized as

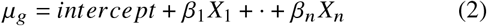

where the values of the dependent factor *μ_g_* are the average coefficient of friction during the displacement step of the finite element simulation.

## Results

We run a total of 38 analyses (26 with full thickness and 12 with the reduced thickness) using FeBio 2.8.2 (MRL The University of Utah, MBL Columbia University) [86]. The postpro-cesing stage is carried out in PostView 2.3 (MRL The University of Utah, MBL Columbia University).

### Strain contours during indentation are sensitive to SC mechanical properties

Representative results from the indentation analysis are illustrated in Figure 3, where the *E*_33_ component of the Green Lagrange strain is plotted for a cross section. The figure shows the results for both dry and wet SC conditions and for three different ages. The flat control is also shown. The maximum magnitudes in *E*_33_ occur in the dermis region under all conditions. The contours of the normal strain resemble Hertzian theory; however, even in the flat control, the different mechanical properties across skin layers contribute to the distortion of the strain contours. There is also an expected boundary effect from the fixed displacements imposed at the bottom of the model. The greatest normal strains occur near the epidermis-dermis interface. The effect of the SC properties is noticeable. Recall that in our model, the dry SC is much stiffer than the wet SC. Accordingly, the strains in the dry SC are smaller and, instead, the deformation is distributed in the other skin layers resulting in extended contours of *E*_33_ in the transverse section. Nevertheless, the magnitude between the two cases remains similar. The contours of other quantities derived from the Green-Lagrange strain (*E*_1_, *E*_11_ and *E*_13_) are available in the supplementary material (Figure S1). These contours further illustrate the notion that the rigidity of the SC influences how the layers participate in carrying the deformation from the indentation. The consideration of the complex surface topography induces a noticeable perturbation of the *E*_33_ contours with respect to the flat geometry. While there is no apparent trend in the contours of *E*_33_ with aging, it is evident that the presence of microrelief leads to asymmetric strain contours inside the tissue, in contrast to the flat geometry. Furthermore, the strain is localized around the different features of the skin topography.

**Figure 3:**
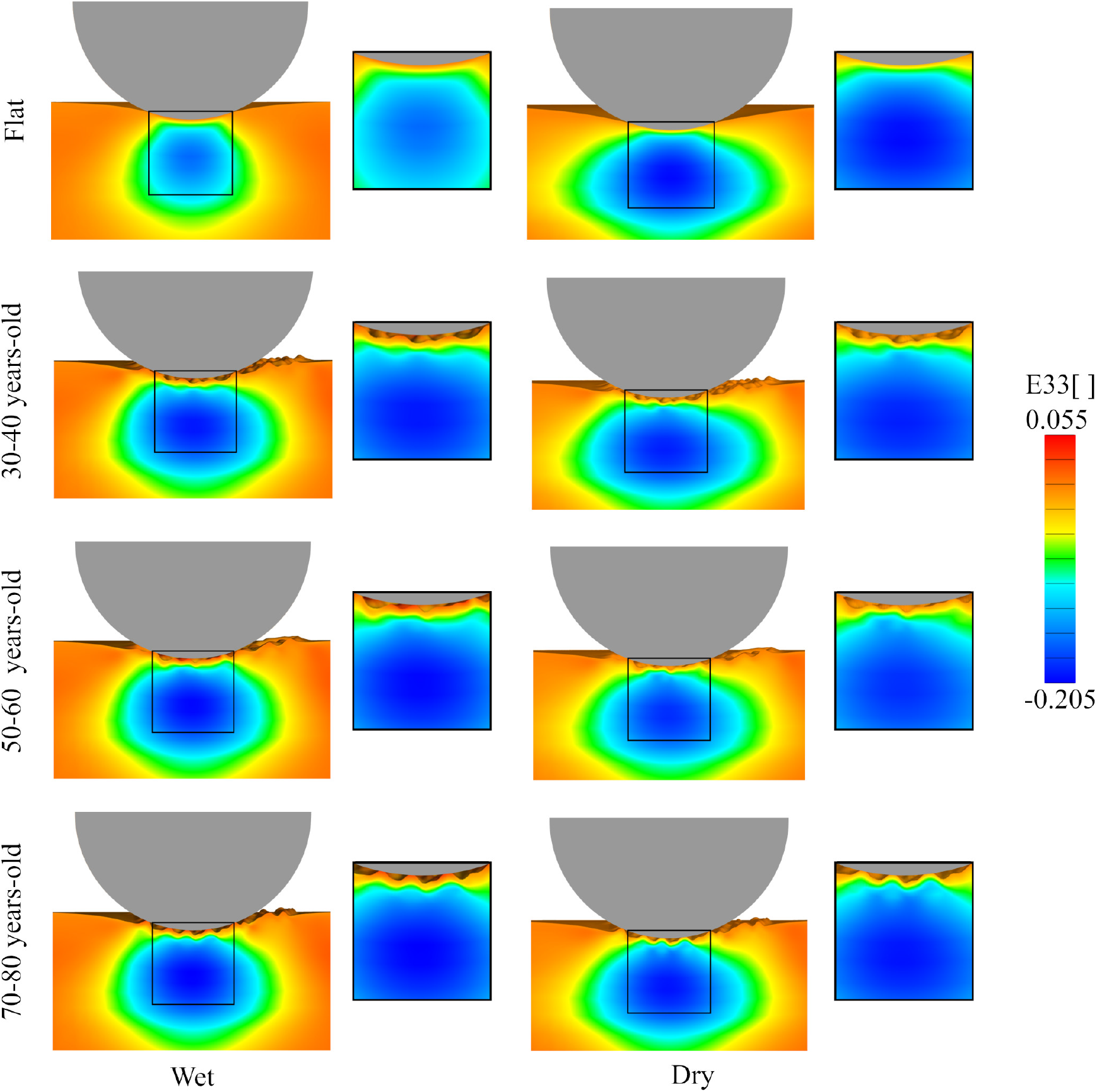
*E*_33_ for different SC condition (wet or dry) for the three different microrelief geometries and the flat control. The strain resembles Hertzian contours. However, even in the flat control, the strain contours are distorted by the changing mechanical properties across skin layers as well as boundary conditions associated with the finite thickness of the skin. The wetness of the SC has a noticeable effect. As the SC dries, its stiffness increases, and the corresponding contours extend over a larger region in the underlying skin layers. The microrelief further contributes to the distortion of the strain contours compared to the flat geometry. In particular, the contours become asymmetric and localized around the features of the surface topography

In Figure 4a and 4b,, both the *E*_33_ and *E*_13_ components of the Green Lagrange strain are plotted along the skin thickness for dry or wet SC conditions and for the different geometries. The plots correspond to elements directly below the center of the indenter. The normal strains are very similar in all cases, whereas the shear strain profiles show significant variation across simulations. The greatest differences in the shear strain between different cases occurs near the top layers of the skin, with *E*_13_ decaying to zero through the dermis and the hypodermis. These variations are related to the surface topography since, as can be seen in Figure 4a, the shear strain in the flat surfaces is small and does not show oscillation in neither the wet nor the dry SC condition. For the detailed geometries, shear strains tend to reach higher magnitude in the wet SC cases compared to simulations with dry SC properties, although this is not always the case. It should also be noted that, even though the microrelief induces shear strains in the skin, these strains are an order of magnitude lower compared to the normal strains depicted in Figure 4b. The normal strain *E*_33_ increases rapidly in magnitude in the epidermis, and then continues to increase in magnitude over the dermis, before going smoothly back to zero over the hypodermis. The plot is consistent in all cases, with small variations with no evident trend.

**Figure 4:**
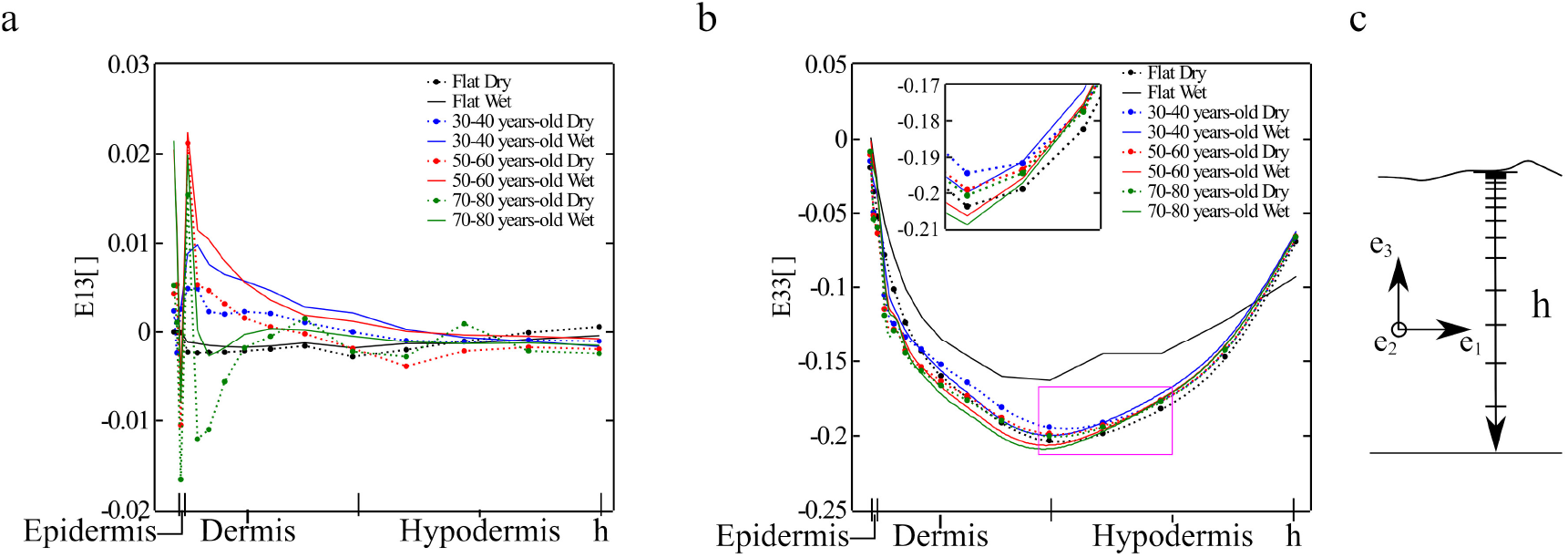
Figure 4: Components of the Green-Lagrange strain plotted across the skin thickness right underneath the indenter for the different age topographies and different SC condition (wet or dry): **a**)*E*_13_,**b**)*E*_33_. The coordinate system is shown in **c**

Therefore, Figures 3 and 4 together indicate that strains in the skin during indentation are mostly normal rather than shear and that the strain profile resembles a Hertzian contour. This is a consequence of the sliding contact interface. However, we also see that the different stiffness across skin layers dictate the deformation across the thickness, with a sharp increase in compression in the epidermis and a maximum compression in the dermis. These observations highlight importance of considering individual skin layer properties when understanding this tissue’s mechanical behavior.

### Influence of age-related changes in the dermis mechanical properties and thickness

Figure 5a shows the *E*_33_ Green-Lagrange strain component for the two oldest microrelief geometries, with a reduced thickness of 50 % with respect to the original model, as well as three different shear modulus values for the dermis. Only the dry SC condition is shown. The change in thickness produces less elliptical and more cone-like shape of the *E*_33_ contours in comparison with Figure 3. Small variations with respect to the shear modulus are observed. To better visualize the change in strain across the thickness and also at different spatial locations with respect to the indenter, components *E*_33_ and *E*_13_ for the 50-60 years-old model are shown in Figure 5b. For each position,the normal strains *E*_33_ have very similar behavior for the three values of the dermis shear modulus, whit a minor differences in the epidermis and dermis zones. The trends also align with the simulations depicted in Figure 4. The shear strain profiles do show significant variation. Right below the center of the indenter the shear strains are small, but they increase as the location of interest is farther from the center and toward the end of the indented region. Additionally, shear strains are larger for the softer dermis.

**Figure 5:**
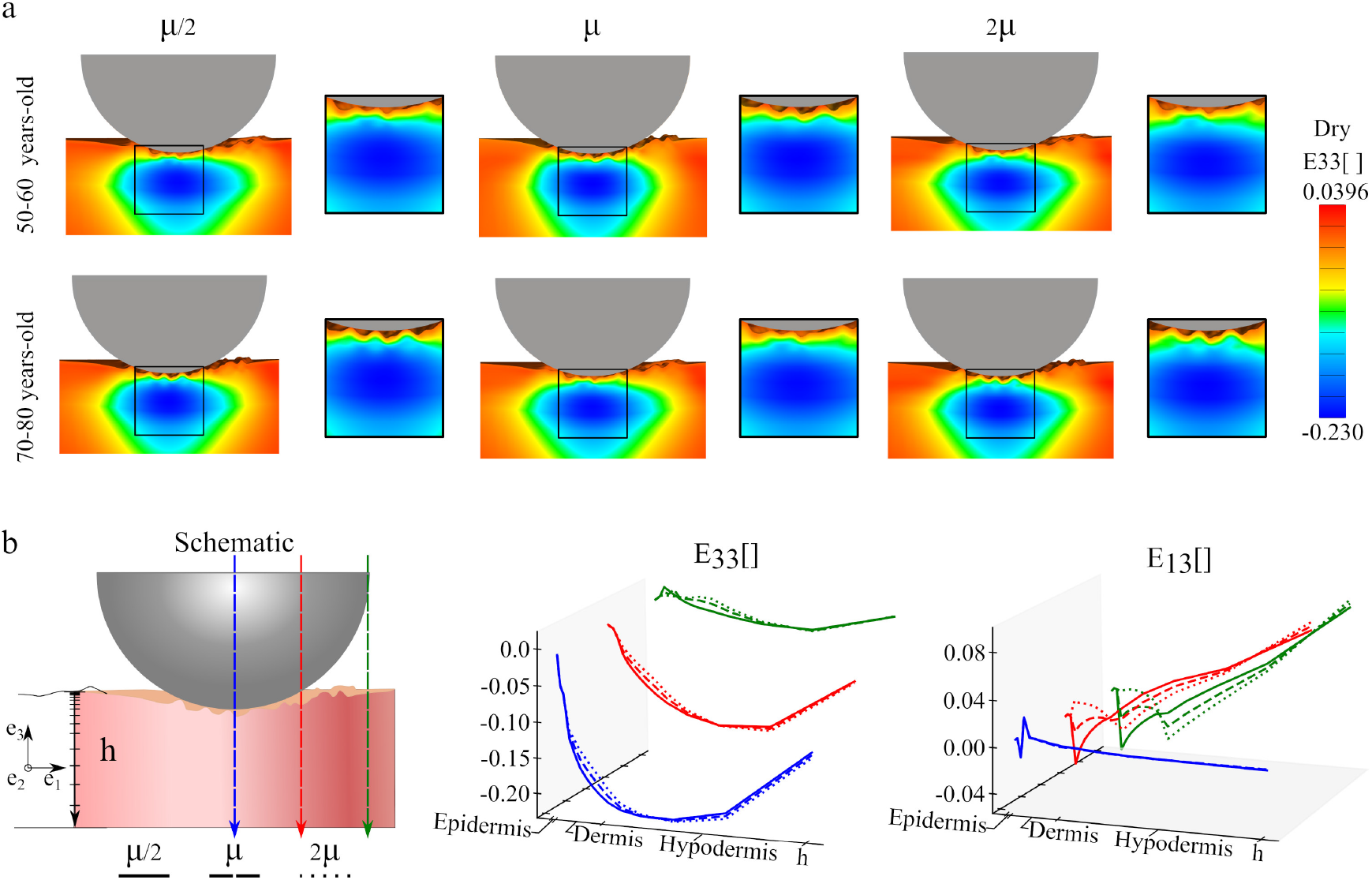
Contours of the Green Lagrange strain components *E*_33_ and *E*_13_ for the two oldest microrelief geometries (reduced thickness model), dry SC condition, and for three values of the shear modulus for the dermis (**a**). *E*_33_ and *E*_13_ plotted across the skin thickness in three different positions underneath the indenter (**b**).

### Influence of aging topography on the surface stress during indenter movement

Following the indentation step, the indenter is displaced along one of the directions shown in Figure 2. First, we focus on the displacement along *U*_1_ and compare the effect of SC properties and the influence of changing geometry with aging. Figure 6 shows the contours of maximum principal stress and maximum shear at the top surface for the three different aging geometries as well as the flat control, and for the two SC conditions. The maximum stress values remain similar for the three different aging topographies, but change noticeably with the change between dry and wet conditions. The changes in principal stress contours for the flat geometry are striking. For the wet case, the SC stiffness is even lower than the underlying epidermis, and the principal stress at the surface of the indented region is negligible. In fact, the greatest principal stress at the surface occurs towards the edge of the indented region. As the SC becomes stiffer, the principal stress contour achieves the maximum at the center of the indented region and decreases away from the indenter. This feature is not observed in the geometries with microrelief.

**Figure 6:**
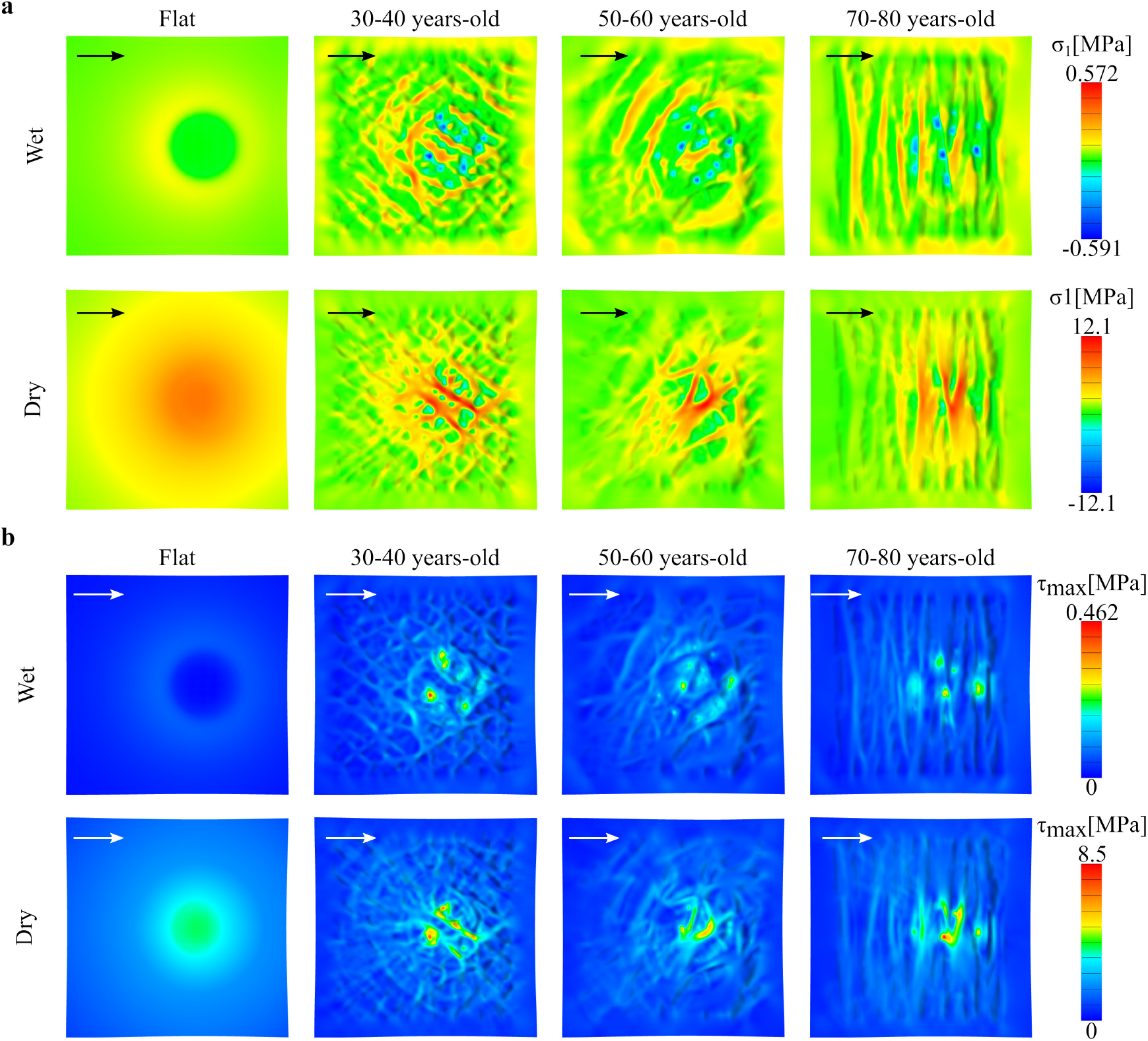
Stress contours at the skin surface for the flat control and the three different skin topographies, for wet and dry SC conditions. The contours correspond to the final position of the indenter after it has moved in the direction *U*_1_ for 1mm at an indentation depth of 500 *μ*m: **a**)Maximum principal stress and **b**) Maximum shear stress.

In the realistic geometries with microrelief, the maximum value of the stress remains relatively unchanged with aging for a given SC condition, with wet SC leading to smaller stress at the surface compared to the dry SC, as expected. However, the most relevant results from the simulations with realistic topography are the patterns of stress distribution over the top surface of the skin due to aging-associated geometric changes. In all cases, the stress is greater in the peaks and plateaus which are in contact with the indenter, and less in the valleys defined by primary and secondary lines, which are not in direct contact with the indenter. Observations of the geometric changes in skin topography with aging translate directly into the observed stress profiles. The primary and secondary lines become more anisotropic with aging, and so do the regions with the peak stresses. For instance, in the 30-40 years-old geometry there is a weaving pattern of high *σ*_1_ stresses, while in the 70-80 years-old geometry there are parallel bands of peak *σ*_1_ stresses. The changes in maximum shear stress are less evident compare to the maximum principal stress.

Table 2 lists the average contact area during the simulation for the flat control surface and the three different skin topographies for the case in which the indenter moves in the *U*_1_ direction. There are a significant difference between SC conditions, reaching more than five times the contact area under wet conditions in comparison to the dry SC case. Clearly, the softer SC in the wet case deforms more and allows for more contact area, 75% larger than dry conditions. The flat control surface has the largest contact area, as expected. The lack of asperities in this scenario allow for an ideal contact between the indenter and skin. The contact area does show changes with aging. As seen in Table 2, contact area increases with aging in the microrelief geometries.

**Table 2:** Average Contact Area [*mm*^2^] for the simulation with the flat skin surface and the three different skin topographies when the indenter moves in the *U*_1_ direction.

| Topography | Contact Area [$mm^2$] | |
| --- | --- | --- |
|  | Wet | Dry |
| Flat | 9.577 e-2 | 2.533 e-2 |
| 30-40 year-old | 7.359 e-3 | 1.334 e-3 |
| 50-60 year-old | 1.129 e-2 | 2.317 e-3 |
| 70-80 year-old | 1.189 e-2 | 2.863 e-3 |

Figure 7 further helps dissect differences in skin topography when the indenter moves in the directions defined by the primary skin lines. The main difference in the magnitude of the stress is still determined by the different rigidity between dry or wet SC. Then, changes in primary skin lines further contribute to distinct patterns of stress over the skin surface, especially in the dry SC case. Due to the symmetry of the primary skin lines with respect the *x*-axis, there are almost no differences between *U*_3_ and *U*_4_. Note, however, that the lines do change with aging, showing increase alignment in the distal-proximal axis (vertical axis in the Figure 7). The reorientation of the primary skin lines, in turn, leads to bands of stress instead of a concentric pattern. This feature of the stress contour was already pointed out for the movement of the indented in the anatomical axis corresponding to the proximal-distal direction *U*_1_, and is further observed in the cases in which the indented moves perpendicular to the skin lines as seen in Figure 7. The stress contours for the indenter moving in the transverse axis *U*_2_ are shown in the supplement (Figure S2). The main insight from moving the indenter in different directions is that, due to increased anisotropy of the primary skin lines, the stress contours also become anisotropic, and this effect is more pronounced in the dry SC case.

**Figure 7:**
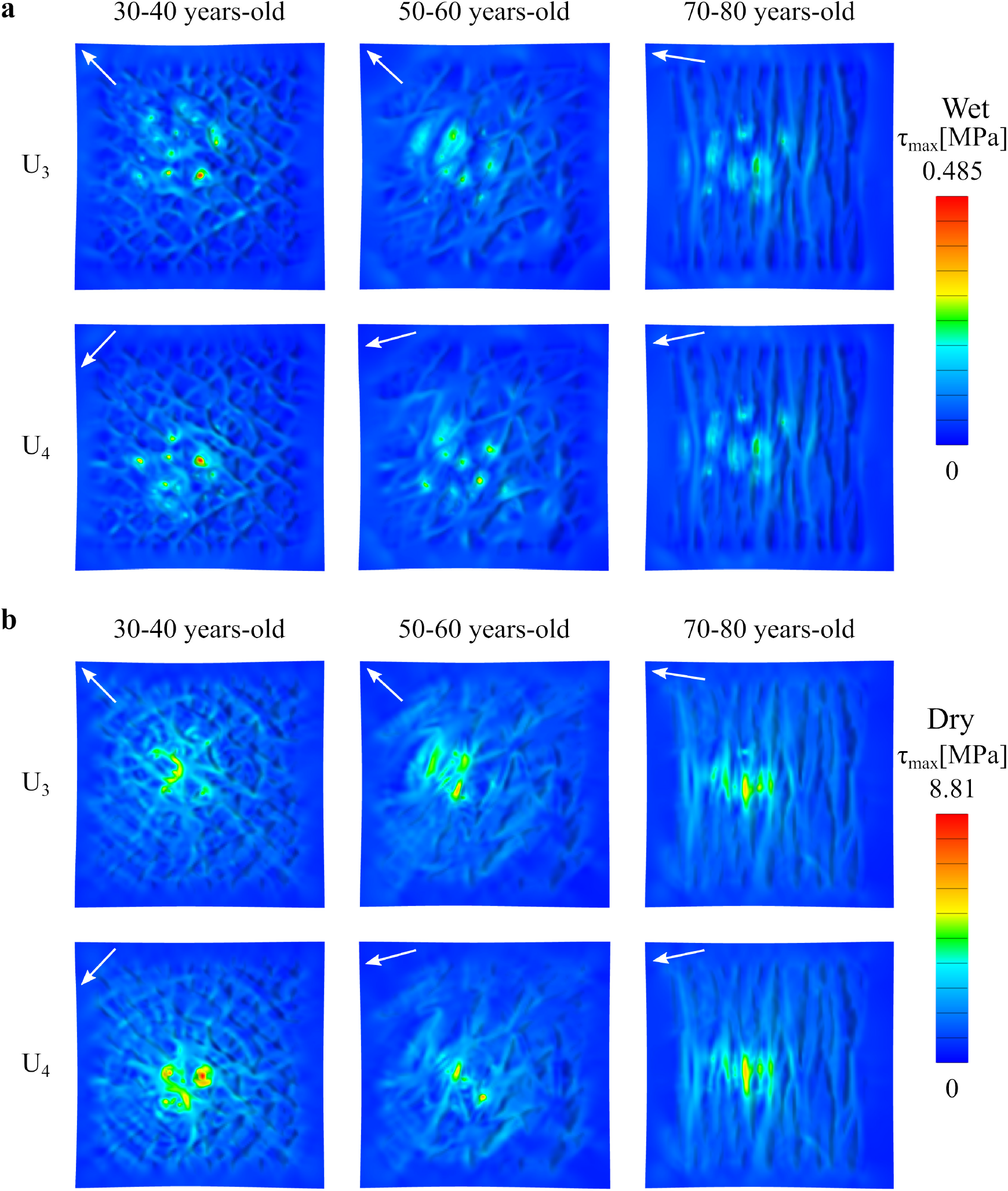
Maximum shear stress contours for three different skin age topographies, for the displacement directions *U*_3_ and *U*_4_ given by the primary skin lines. The results correspond to: **a**)wet SC and **b**)dry SC material properties

### Variation in the reaction forces as a function of aging microrelief

Accompanying the changes in stress distributions, the topographical changes of the skin with aging are also reflected in the reaction forces during indenter movement, which are plotted in Figure 8. In Figure 8, *F_t_* is the component of the reaction force in the direction of the indenter movement, *F_n_* is the normal reaction force, i.e. normal to the horizontal plane, while *F_b_* is the force component orthogonal to the other two directions of interest. In other words, *F_b_* is also in the horizontal plane, but orthogonal to *F_t_*.

**Figure 8:**
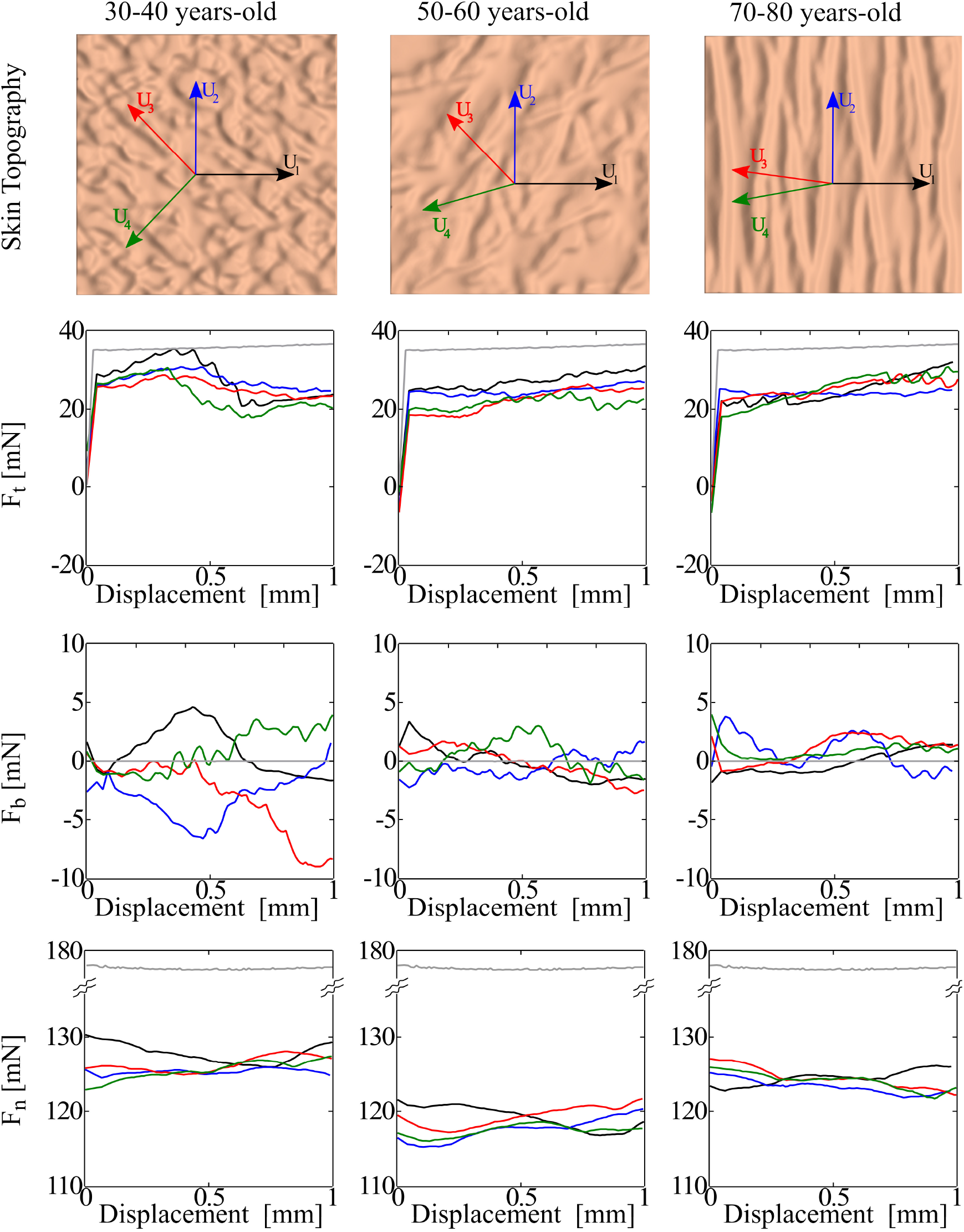
Behavior of in-plane (*F_t_* and *F_b_*), and normal (*F_n_*) reaction forces on the indenter for three different skin age topographies and four displacement directions (black: *U*_1_, red *U*_2_, blue *U*_3_, green *U*_4_). The forces corresponding to the flat control are shown in grey. All plots correspond to the properties of the dry SC

The normal component of the force is very similar in all cases, oscillating between 115 and 130 mN. In comparison, the normal force corresponding to the flat geometry (grey line in Figure 8), is nearly constant at 175 mN. For context, the coefficient of friction has been assessed in experiments with normal forces ranging between 5mN and 8N [87, 88]. The normal forces obtained in this study are in the ranges of normal forces as reported experimentally [43, 30, 54]. The tangential force for the flat geometry is also nearly constant during indenter movement, at *F_t_* = 35 mN. For the realistic geometries, the tangential force directly opposing the indenter movement oscillates around 20 mN. Interestingly, the in-plane component F¿ is non-zero in the realistic geometries. In the control case of the flat geometry *F_t_*, =0 as expected. In the geometries with microrelief, the force *F_b_* oscillates between −10 and 10 mN. We remark that these forces orthogonal to the expander movement appear because we fully control the displacement of the indenter, such that the deformation of the microrelief as the indenter moves in a given direction can be asymmetric with respect to the plane defined by the indenter movement. From the plots in Figure 8, there is no notable difference between the different direction of indenter movement. Aging does contribute to observable trends. Changes of the microrelief with aging tend to reduce the oscillation of the tangential forces. This can be explained by the surface features with aging. In the 30-40 case, the primary lines form a weaving pattern such that the indenter crosses many obstacles as it slides on the skin. With aging, as the primary skin lines become more anisotropic there is less variation in the obstacles that the indenter has to go over.

### Strain contours during indenter displacement

Figure 9 shows the *E*_11_ contours as the indenter moves, further clarifying the trends in the reaction forces. In the flat geometry, the E⊓ contours are unchanged as the indenter moves, which is expected and aligns with the nearly constant *F_t_* in Figure8 for the flat geometry. We remark that the E⊓ strain corresponds to the normal strain in the direction of the indenter movement. Therefore, this component of the strain can contribute to the tangential force *F_t_* in addition to the friction force due to local contact between skin and indenter. Indeed, while the friction force between the indenter and the skin surface is a primary contributor for *F_t_*, previous work by Tang et al, Derler et al and Leyva-Mendivil et al, on contact between indenters and soft substrates identified a deformation component of *F_t_* [31, 34, 89]. This deformation component is characterized by a bulging ahead of the indenter as it moves. In our case, it does not appear that the skin deforms asymmetrically in the flat geometries. Consequently, the *E*_11_ contours appear to be symmetric with respect to the indenter movement direction. Calculating the global friction coefficient based on the *F_t_* and *F_n_* forces reported in Figure8 leads to *μ_g_* = 0.2043 under dry conditions (*μ_g_* =0.2009 on wet conditions), which is very close to the local friction coefficient *μ_l_* =0.2. This confirms the observation that the *E*_11_ contours are mostly symmetric with respect to the indenter movement and that there is a negligible deformation component for *F_t_* in the flat geometries. The contours of *E*_11_ in the geometries with microrelief can help explain the variation in *F_t_* observed in Figure8. The overall strain contours are similar in shape and magnitude to each other, but the microrelief has two evident effects. First, the E⊓ contours are no longer constant as the indenter moves, but change due to the surface features of skin. Secondly, the contours are no longer symmetric with respect to the plane determined by the indenter movement direction. It can be seen in the zoom-in panels, that the strains are greater in the peaks which are in contact with the indenter. Thus, for the realistic geometries, the variation in the topography leads to oscillating tangential forces given by both changes in contact area as well as the underlying mesoscale deformation. At the same time, the realistic geometries do not show a bulging ahead the indenter, which would be reflected in the *E*_1_1 contour as a clear asymmetry. Thus, we do not expect a significant deformation contribution to *F_t_*. This is further discussed in the next section.

**Figure 9:**
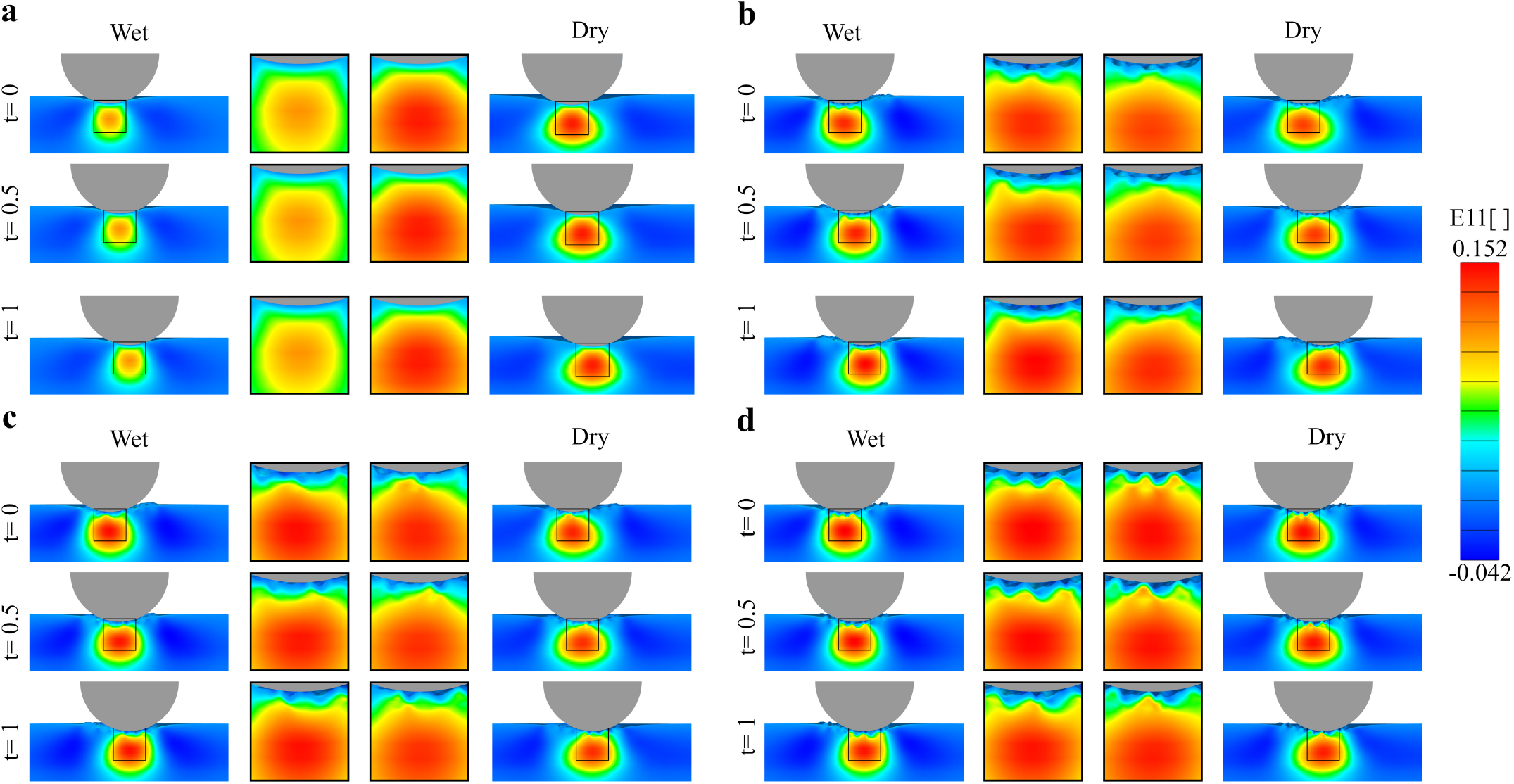
Normal strain *E*_11_ in the direction of the indenter movement at three time points during indenter displacement (*t* = 0, 0.5, 1s), for the two SC conditions, and for the four different geometries:**a**)Flat, **b**)30-40 years-old, **c**)50-60 years-old and **d**)70-80 years-old

### Global coefficient of friction

Based on the values for *F_t_, F_t_* and *F_n_* for all simulations, we calculate the global coefficient of friction according to eq. 2. Table 3 summarizes the results. The global coefficient of friction in our simulations with microrelief is in the range *μ_g_* ∈ [0.1845,0.2306]. For the flat geometry, global coefficient of friction is *μ_g_* ∈ [0.2009,0.2043] for wet to dry SC. The local coefficient of friction is *μ*_11_ = 0.2. The range of *μ_g_* for the simulations with microrelief suggests that the topography of skin contributes to the frictional response. In general the dry SC has higher values than wet state with some exceptions. This results are associated to the deformation of the ridges in wet state reducing the ploughing effect of the skin on the indenter..

**Table 3:**
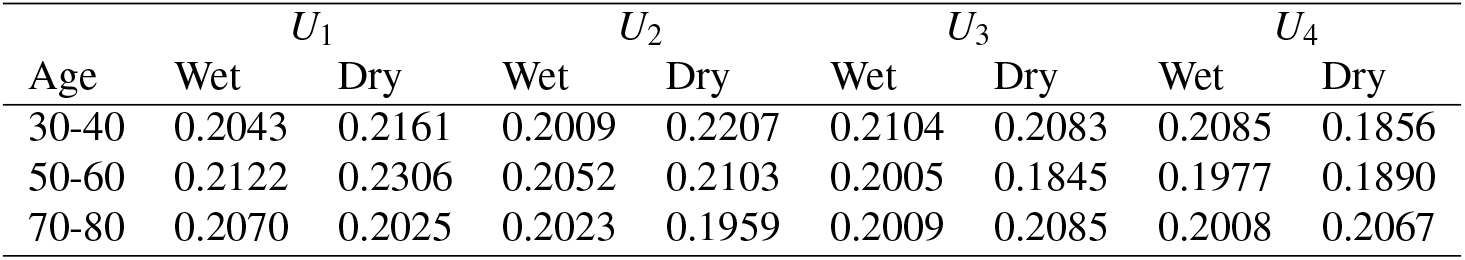
Coefficient of friction for three different skin topographies, four displacement directions, and two conditions of the SC

Based on the result of the multivariate regression analysis with the generalized linear model for the coefficient of friction, we describe the coefficient of friction through

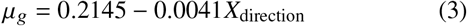

In this model, it is considered that the coefficient of friction follows a Gaussian distribution. The model has an adjusted generalized R–squared value of *R_adj_*=0.16, Ramsey’s Reset for Specification for the model Reset=1.0183 with a p-value=0.3792, and the residuals distribute normal according to a Jaque-Bera test has p-value=0.9755 (see Table S1 in the supplementary material). According to the regression analysis only the displacement direction of the indenter has significant influence in the coefficient of friction (p<0.05). The displacement direction correlates negatively with the coefficient of friction, increasing the codification value (1, 2, 3 and 4 for *U*_1_, *U*_2_, *U*_3_ and *U*_4_ respectively) the coefficient of friction is reduced, suggesting lower friction against the primary lines. The other parameters have lower or null contributions in the phenomena under the conditions analyzed in this paper.

## Discussion

Understanding the frictional behavior of skin is key for the design of new technologies, from sportswear and wearable gadgets, to medical devices. The interaction between skin and other materials is complex and, due to inherent limitations with experimental setups, it is challenging to resolve the effect of the skin’s microrelief deformation on the measured frictional forces. Numerical investigations have offered some clues [33, 34, 45]. However, while it is known that the topography of skin changes with age, the effect of this change has not been investigated. Moreover, only two-dimensional analyses have been [33, 34]. Yet, the topography of skin has features in three dimensions. To address these gaps, here we show a numerical investigation of the skin interacting with a rigid indenter accounting for the 3D features of the skin microrelief and their change with aging.

### Strain contours under indentation depend on individual skin layer properties

The indentation simulations showed that the topography has a small effect on the strain contours in the skin. We recovered contours resembling classical theory of elastic contact [4]. We further confirmed the shape of the contours in idealized geometries without any microrelief. The most important factor influencing the deformation of skin under the indenter was the presence of individual layers with different mechanical properties. Indeed, the need to consider individual skin layers has been advocated by many, particularly for the type of loading present during indentation [1, 2, 76, 90, 91]. Furthermore, it has been reported in the literature that the change in humidity has a marked effect on the mechanical properties of the outermost layer, the SC. Our simulations show that the stiffening of the SC with decrease in humidity leads to a change in the distribution of the strains over a larger region, but the magnitude of the strains remains relatively unchanged. It remains to test whether other static loading scenarios relevant to the skin physiology would be more visibly affected by the change in the SC properties, or if the change in strain distribution in the skin from changes in SC mechanics has secondary effects linked to skin disease. For instance, pressure ulcers form when the skin is subject to compressive loading, particularly near bony prominences [7, 9, 16, 92, 93]. Our work in developing a detailed 3D model of skin including individual layer properties can therefore be useful in the study of pressure ulcer mechanics or other scenarios with similar loading on skin.

We also looked at changes of mechanical properties in the dermis as well as thickness changes associated with aging. There was little effect on the strain contours and friction coefficients. This is because the simulation is kinematically driven. However, it would be interesting to investigate the response under prescribed loading, which might be more appropriate in a clinical setting.

### Microrelief aging leads to anisotropic stress patterns during skin-indenter contact

Previous reports have quantified the changes in skin topography or microrelief with aging as well as changes in friction [44, 46, 50, 94, 95, 96]. We were therefore poised to evaluate if some of those changes could be explained in our simulation with using different 3D geometries of microrelief. Indeed, we see that the microrelief becomes more anisotropic with aging, with the primary skin lines becoming more aligned in the proximal-distal direction in the forearm. Consequently, movement of the indenter lead to more anisotropic stress patterns. Since we use the same material properties the only change we saw was due to the geometry. We did see some changes in stress contours as we changed the mechanical properties based on humidity. Wet or dry SC properties mainly preserved the patterns but altered the magnitude of the shear and normal stresses at the SC. These simulations therefore imply that in designing surfaces in contact with skin it is important to consider the skin topography in three dimensions, as aging will lead to anisotropic stress distributions aligned with the primary skin lines. A related work recently showed that the microrelief can also explain buckling and furrows of the skin [45]. However, the work in [45] considered a single adult geometry. Our work therefore also opens new questions, such as the contribution of aging microrelief to buckling under compression. As mentioned before, the surface geometry is not the only thing that changes with aging, but also thinning of the dermis [44, 97, 84], and changes in mechanical properties [44, 47, 98, 99, 100]. We did preliminary work in this direction and we will continue to investigate these changes in more clinically relevant cases in the near future.

In addition to the stress patterns becoming more anisotropic, we also found that the contact area is highly affected by the changes in surface topography with aging. With aging, there is an increase of the plateaus and a reduction of the number of furrows on the surface [57, 58]. Thus, with larger plateau area the contact with the indenter increases in older microrelief geometries.

### The primary skin lines in the microrelief influence the global coefficient of friction

This work focused on the interaction with the skin using a local friction coefficient *μ_l_* =0.2. Previous experimental studies have shown that the interaction of the human skin with different materials and surfaces can show coefficient of friction over a wide range, between 0.059 and 3.7 [101, 43, 102]. Although our analysis does not represent the interaction between the human skin and particular materials, values of *μ*=0.25 have been reported for the interaction between calf skin and nitrocellulose [29], *μ*=0.22 for the contact between forearm skin and stainless steel [103], *μ*=0.22 between forearm and polypropylene and between different body zones and Teflon [27], and *μ*=0.19 between forearm skin and steel [43] among other similar values to the local coefficient of friction considered here. Previous work considering rigid indenters and soft substrate showed that the reaction forces on the indenter are not just due to the local friction but that an asymmetric deformation field on the length scale of the indenter can contribute a significant deformation component [33, 34, 85]. In the presence of stronger adhesion interactions, skin at the front of the indenter is compressed while tissue gets stretched behind the indenter movement [104]. Different ranges of material properties, indentation depth, local friction coefficient, and adhesion are needed to see a significant deformation component, which remains to be studied. In our case, keeping all the parameters the same and changing solely from flat to the realistic microrelief surfaces we see an effect in the coefficient of friction [1]. Looking at the strain contours in a transverse cross section we see that the strains change as the indenter moves on top of the skin and that the contours are asymmetric with respect to the plane defined by indenter movement. However, the features in the strain contours are on the mesoscale, dictated by the ridges and valleys of the microrelief, and while contribute to the overall oscillation of the tangential forces, there is no significant contribution to a deformation component opposing indenter movement. As a result, the average friction coefficient in the realistic geometries is close to the local friction coefficient, within 13 %. In Previous 2D studies by Leyva-Medivil et. al., smaller indenters, with size similar to the topographical features, resulted in a noticeable deformation component [33]. To understand the contribution between deformation and adhesion to the overall frictional response, it is useful to look at the ratio between the skin surface features and the indenter surface features. Consider the ratio *ψ = φ/R_a_* or *ψ = φ/R_t_*, where *φ* is the diameter of a perfectly smooth indenter, *R_a_* is the arithmetic average height of the skin topography, and *R_t_* is the maximum height of the skin topography (e.g. distance between the deepest valley and the highest peak) [105]. *R_a_* for human skin is about 6.7 *μ*m – 46.2 *μ*m and *R_t_* is between 74.03 *μ*m and 84.3 *μ*m [30, 54, 95, 106, 107]. Numerical studies performed by Leyva-Mendivil et al [33] considered *ψ* between 0.8 and 4 (base on *R_t_*) and showed a notable influence of the deformation component in the frictional response. In our case *ψ* is above 31 (base on *R_t_*), which helps to explain the small role of the microrelief deformation on the global friction coefficient.

The more interesting trend of the multivariate analysis was the effect of direction. This was expected form the stress analysis, where we saw more anisotropic stress patterns due to the reorientation of skin lines with aging. Nevertheless, the changes are small, especially considering possible variation across individuals. Clearly more work is needed, including adding an adhesion component, testing more indenters or surfaces with asperities, different local friction coefficients, and a larger set of geometries and boundary conditions.

### Limitations

While this work explores the effect of aging in 3D geometries on friction for the first time, it still could improve from a wider set of conditions. For instance, this study focuses on the skin of the forearm. Skin is classified into glabrous and hairy: glabrous skin is that of the palms and soles of the feet and hairy skin covers the rest of the body [108]. Our simulations consider the forearm skin topography but do not include the presence of hair. Thus, it remains to look at other anatomical regions with different skin morphology, topography, mechanical properties, and hair bearing characteristics, such as buttocks or amputation stumps. Mechanical properties of human skin layers were considered as isotropic and from a few sources [76, 45, 33]. More sophisticated material models and more data is also needed in this regard. For instance, several microstructure-based models that account for collagen and elastin fibers and therefore tissue anisotropy have been proposed and should be considered [37]. We ignored adhesion and considered a single local coefficient of friction. Again, a broader parameter range driven by literature reports is needed to include additional variables in a larger study. A single indenter geometry and indentation depth were considered, but other geometries and indentation depths are certainly expected to improve our understanding of skin friction. Lastly, we only considered three microrelief geometries. While this is clearly an improvement compared to simplified 2D geometries or 3D models of skin that ignore microrelief, additional 3D geometries with microrelief are certainly needed to obtain statistically valid results. At the same time, the changes in microrelief that we see in these three geometries are reflective of the overall changes of microrelief with aging that have been thoroughly reported [46, 35, 56, 57, 58].

## Conclusions

We show the importance of multi-layered 3D geometries of skin with realistic microrelief and changes with aging. The different material properties across skin layers are essential for both the indentation and movement steps. The patterns of the microrelief in 3D are characterized by more anisotropic primary lines which affect the stress distribution of the skin in contact with the indenter and ultimately affect the friction coefficient depending on orientation. Therefore our results underscore the importance of considering realistic geometries in 3D, especially multi-layered properties and surface topography, for skin mechanics and tribology. This will enable better design of medical devices and textiles particularly accounting for aging effects.

## Acknowledgements

This work was carried out under support of Colombia Ministry of Science, Technology and Innovation MINICIENCIAS (COLCIENCIAS Doctorado Nacional 647 -2014/Universidad de Antioquia). The financial support of the Universidad de Antioquia is also gratefully acknowledged. This work as also partially supported by award CMMI 1916668 to Adrian Buganza Tepole.

