## Supplement for "Changes in the three-dimensional microscale topography of human skin with aging impact its mechanical and tribological behavior"

### Principal strains during skin indentation

The main text shows the normal strains in a cross section of the domain as skin is indented (Figure 3). In the supplement we further show the first principal Green-Lagrange strain $E1$, as well as the components $E_{11}, E_{12}$ to further illustrate the role of microrelief and individual skin layer properties on the strain dsitributions over a central cross-section of the three-dimensional domain. Figure S1 shows that, in the wet condition, for which the top layer of the skin is soft compared to the underlying layer, the epidermis, indenttion of a flat geometry leads to strains which increase gradually over the epidermis and are largest in the dermis, after which they decrease over the hypodermis. Accounting for microrelief realds to higher strains which propagate all the way down to the bottom of the domain. The reason for this increase in strain is that the microrelief geometry consists of peaks and valleys, thus, even though the distance of the indenter to the equivalent flat skin surface remains the same, the microrelief has features above this line which interact with the indenter sooner with respect to the flat case. In the dry condition, characterized by stiffer SC, the strains are higher compared to the wet SC.

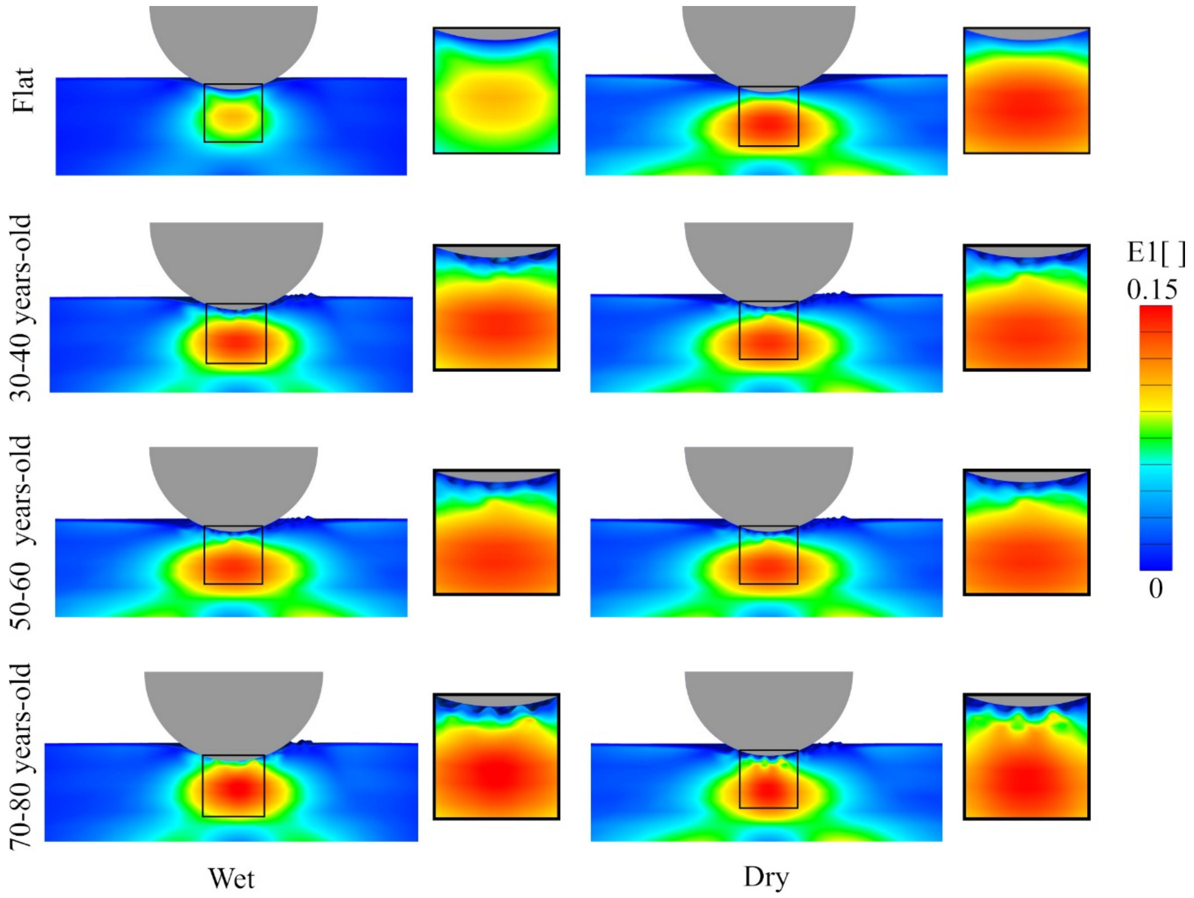

Figure S1 E1 for different SC condition (wet or dry) for the three different microrelief geometries and the flat control.

The contours of the $E_{11}$ component of the Green-Lagrange strain in Figure S2 show much less influence from microrelief changes or changes of SC properties between wet and dry conditions compared to Figure S1 or Figure 3 in the main text. Indeed, the indentation step mostly affects the normal strain component or the first principal strain. Similarly, Figure S3 shows that the shear strain component is minimally affected by the changes in microrelief or changes in SC properties. Nevertheless, Figure S2 does show that the microrelief leads to strain concentration around the microscale features, mostly in the epidermis, which might be important for improving understanding of epidermis mechanosensing.

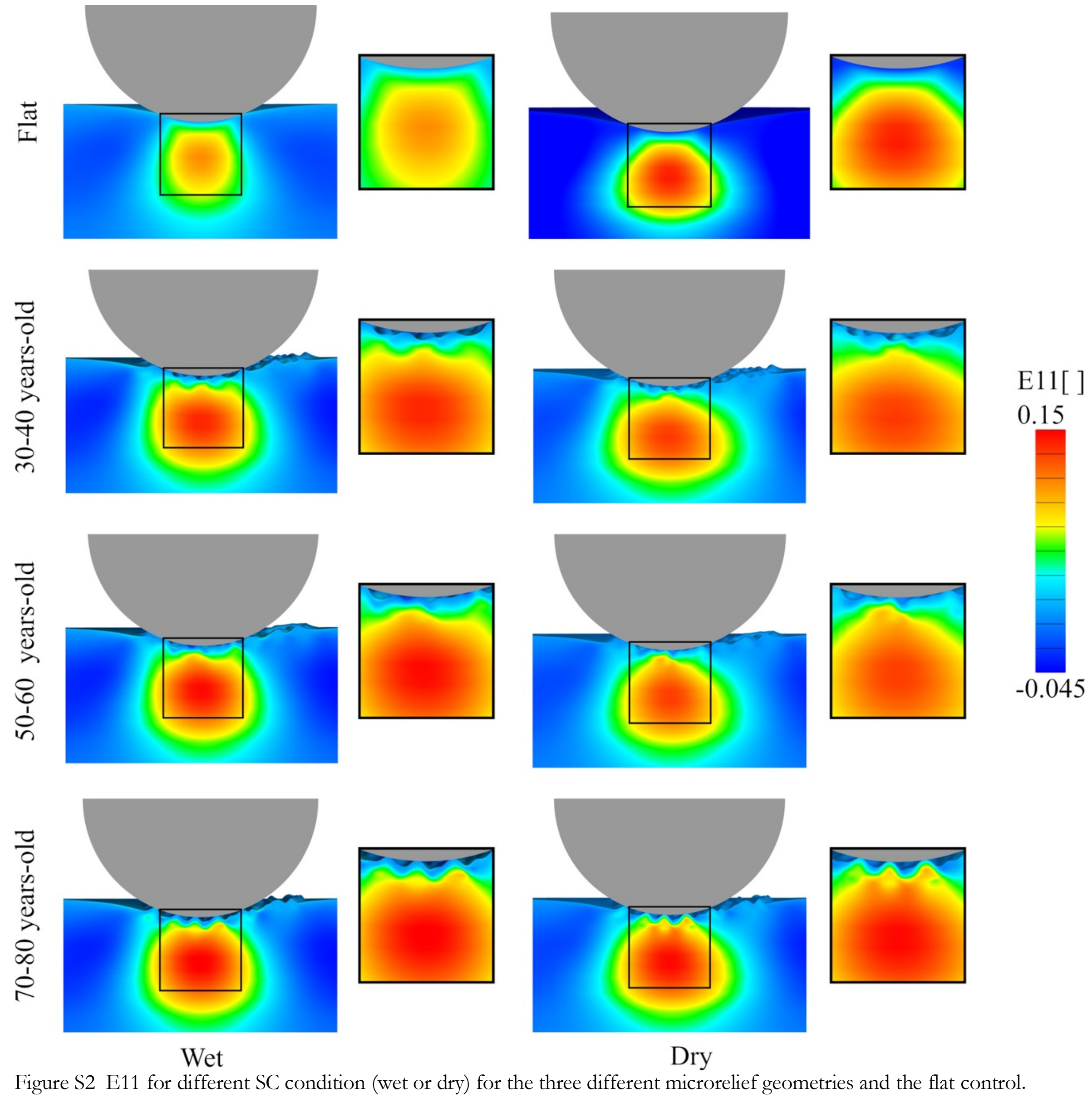


Figure S2 E11 for different SC condition (wet or dry) for the three different microrelief geometries and the flat control.

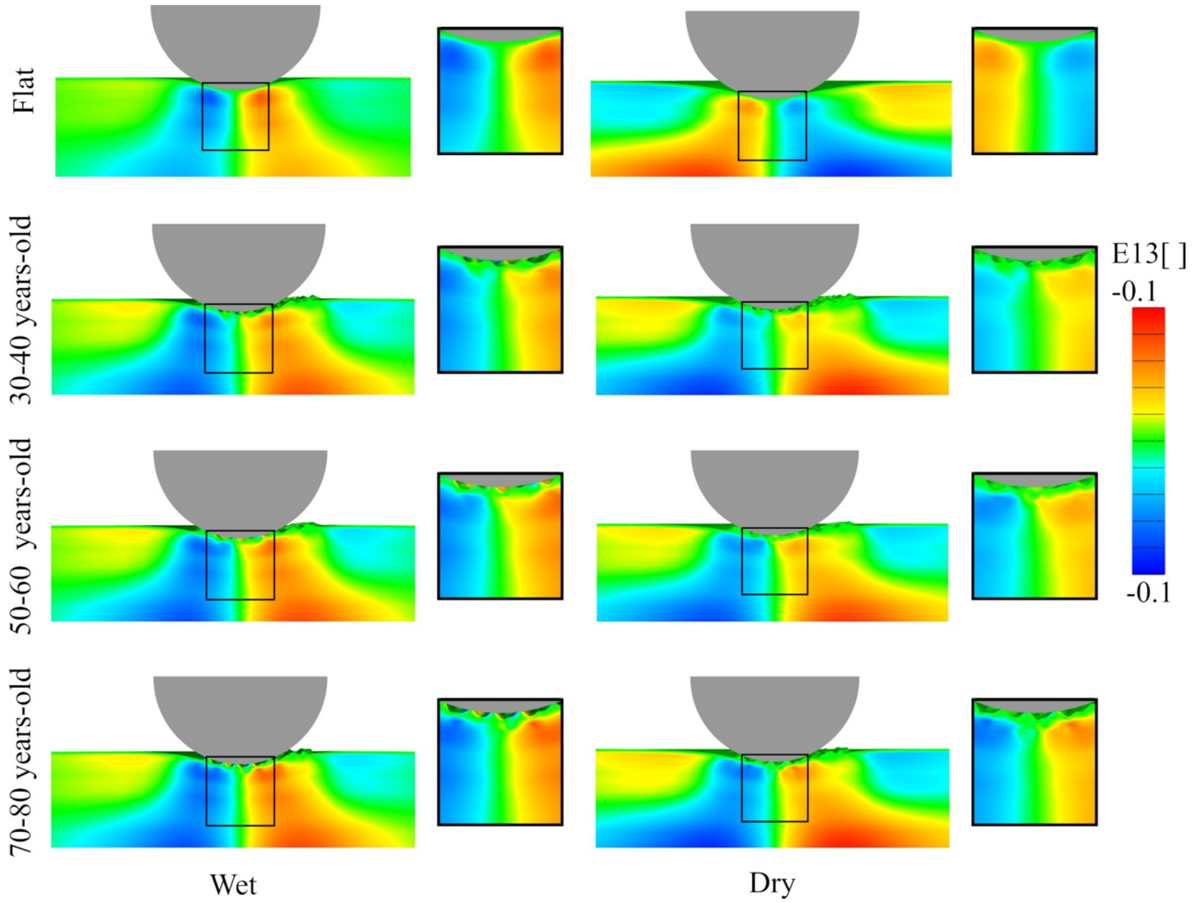

Figure S3 E13 for different SC condition (wet or dry) for the three different microrelief geometries and the flat control.

**Surface stress during indenter movement**

The main text shows the maximum principal stress at the skin surface when the indenter moves in the anatomical directions $U_1$ which corresponds to the proximal-distal axis. Figure S4 shows the maximum principal stress when the indenter moves in the directions given by the primary skin lines $U_3$ and $U_4$. Similar to the discussion in the main text, the skin microrelief becomes more anisotropic with aging, which leads to an increasingly anisotropic stress profile at the surface as the skin interacts with the indenter.

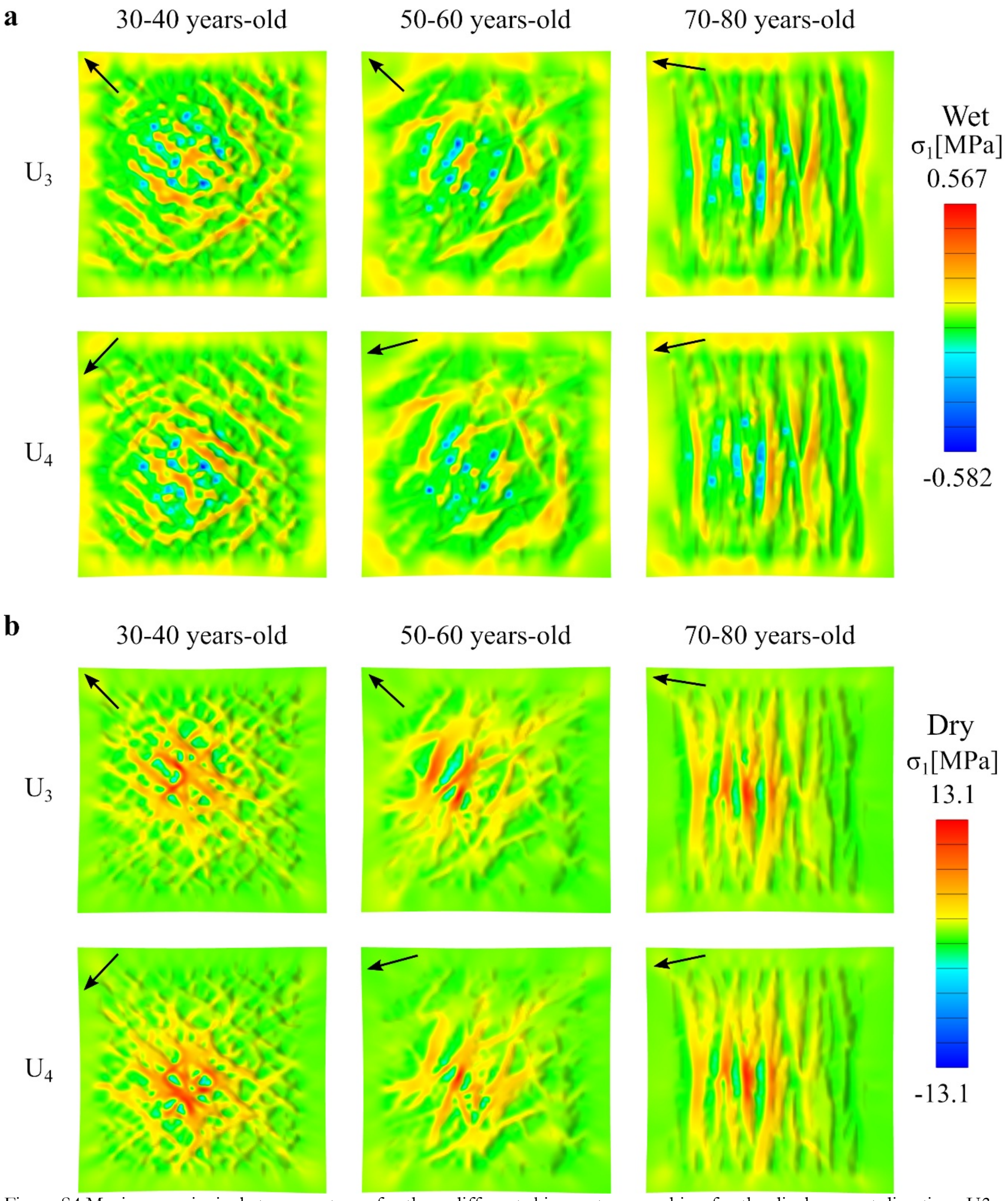


Figure S4 Maximum principal stress contours for three different skin age topographies, for the displacement directions U3 and U4 given by the primary skin lines. The results correspond to: a) wet SC b) dry SC material properties.

### Regression for skin friction as a function of model parameters

As indicated in eq. (2), we perform a simple multi-linear regression of the global coefficient of friction in terms of the parameters of the model. The corresponding R functions are results are illustrated here. As can be seen from the results, only the direction of indenter movement with respect to the skin microrelief lines show statistical significance for the global coefficient of friction, whereas neither age nor SC properties led to statistically significant results. A second linear regression was then performed, ignoring all parameters except for direction. These results are shown in Table S2.

**Table S1 Generalized linear model for all variables**

```
Call:
glm(formula = tableFriction$GCOF ~ tableFriction$Age + tableFriction$SC +
tableFriction$Direction, family = gaussian)

Deviance Residuals:
      Min         1Q     Median         3Q        Max
-0.060141  -0.007372  -0.001449   0.008028   0.069904

Coefficients:
                          Estimate Std. Error t value Pr(>|t|)
(Intercept)              0.2107849  0.0011905 177.050   <2e-16 ***
tableFriction$Age        0.0001986  0.0003715   0.535    0.593
tableFriction$SC         0.0010778  0.0007171   1.503    0.133
tableFriction$Direction -0.0032078  0.0003206 -10.006   <2e-16 ***
---
Signif. codes:  0 ‘***’ 0.001 ‘**’ 0.01 ‘*’ 0.05 ‘.’ 0.1 ‘ ’ 1

(Dispersion parameter for gaussian family taken to be 0.0003155185)

    Null deviance: 0.81115  on 2468  degrees of freedom
Residual deviance: 0.77775  on 2465  degrees of freedom
AIC: -12891

Number of Fisher Scoring iterations: 2
```

**Table S2 Generalized linear model for Direction**

```
Call:
glm(formula = table$GCOF ~ table$Direction, family = gaussian)

Deviance Residuals:
       Min          1Q      Median          3Q         Max
-0.0177857  -0.0055959  -0.0008867   0.0058259   0.0202161

Coefficients:
                 Estimate Std. Error t value Pr(>|t|)
(Intercept)      0.214449   0.004695   45.68   <2e-16 ***
table$Direction -0.004046   0.001714   -2.36   0.0275 *
---
Signif. codes:  0 ‘***’ 0.001 ‘**’ 0.01 ‘*’ 0.05 ‘.’ 0.1 ‘ ’ 1

(Dispersion parameter for gaussian family taken to be 8.817396e-05)

    Null deviance: 0.0024310  on 23  degrees of freedom
Residual deviance: 0.0019398  on 22  degrees of freedom
AIC: -152.05

Number of Fisher Scoring iterations: 2
```

**Mesh convergence**

To verify that the mesh density was accurate we performed a mesh refinement analysis and found that increasing the mesh by a factor of 1.5 resulted in small variations of the stress features. In particular, the maximum shear stress shown in Figure S5 only increases by 0.8% with the 50% increase in mesh elements. The simulations are already computationally demanding with the *coarse* mesh which consists of approximately 350,000 hex elements.

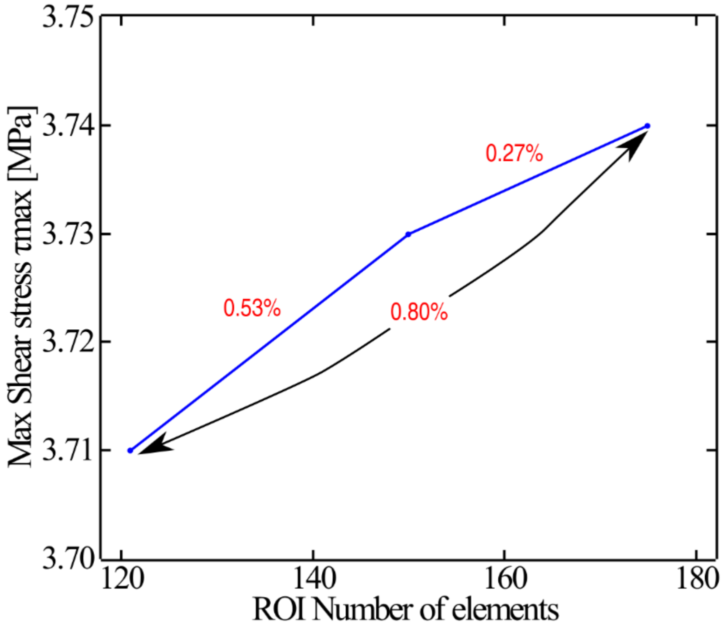


Figure 5 Mesh convergence analysis for Max Shear stress $\tau_{max}$ of skin flat topography. The mesh is based on Preview Butterfly 2D 120,150,175 elements in x and y direction for 3.5 mm square ROI.
